# Short Neurorobotics Workshop for High School Students Promotes Competence and Confidence in Computational Neuroscience

**DOI:** 10.1101/597609

**Authors:** Christopher A. Harris, Stanislav Mircic, Zachary Reining, Marcio Amorim, Đorđe Jović, Lucia Guerri, William Wallace, Gregory J. Gage

## Abstract

Understanding the brain is a fascinating challenge, captivating the scientific community and the public alike. The lack of effective treatment for most brain disorders makes the training of the next generation of neuroscientists, engineers and physicians a key concern. Over the past decade there has been a growing effort to introduce neuroscience in primary and secondary schools, however hands-on laboratories have been limited to anatomical or electrophysiological activities. Modern neuroscience research labs are increasingly using computational tools to model circuits of the brain to understand information processing. Here we introduce the use of neurorobots - robots controlled by computer models of biological brains - as an introduction to computational neuroscience in the K-12 classroom. Neurorobotics has enormous potential as an education technology because it combines multiple activities with clear educational benefits including neuroscience, active learning, and robotics. We describe an introductory neurorobot workshop that teaches high school students how to use neurorobots to investigate key concepts in neuroscience, including spiking neural networks, synaptic plasticity, and adaptive action selection. Our do-it-yourself (DIY) neurorobot uses wheels, a camera, a speaker, and a distance sensor to interact with its environment, and can be built from generic parts costing about $150 in under 4 hrs. Our Neurorobot App visualizes the neurorobot's visual input and brain activity in real-time, and enables students to design new brains and deliver dopamine-like reward signals to reinforce chosen behaviors. We have tested the Neurorobot Workshop with high school students (n = 3 workshops, 9 students total) and have found that students were able to complete all exercises in under 3 hrs. In a post-workshop survey, students reported having gained the ability to develop neural networks that perform specific functions, including goal-directed behavior and memory. Here we provide DIY hardware assembly instructions, discuss our open-source Neurorobot App and demonstrate how to teach the Neurorobot Workshop. By doing this we hope to accelerate research in educational neurorobotics and promote the use of neurorobots to teach computational neuroscience in high school.

## 1. INTRODUCING EDUCATIONAL NEUROROBOTICS

Understanding the brain is necessary to understand ourselves, treat brain disorders and inspire new scientists. Insights from neuroscience are also facilitating rapid progress in artificial intelligence (Hassabis et al., 2017; Zador, 2019). Nevertheless, most students receive almost no education in neuroscience and the public’s understanding of the brain is lacking (Dekker & Jolles, 2015; Fulop & Tanner, 2012; Sperduti et. al., 2012; Labriole 2010; Frantz et al., 2009). The reasons cited are that the brain is perceived to be too complex, and that the tools needed to study it are too expensive and hard to use. Although neuroscience is not yet an independent component of typical high school curricula, some schools are adopting neuroscience courses or steering their biology or psychology classes in the direction of neuroscience to satisfy growing interest in the brain (Gage, 2019).

An important cause of the increasing prominence and appeal of neuroscience in recent years is its powerful synergy with computer technology. Brain imaging and visualization techniques have given neuroscientists and the public unprecedented access to the complex structures and dynamics of brains. Computer modelling is enabling researchers to go beyond theorizing about brain function to actually implementing those functions *in silico*. Large networks of simple neurons connected by plastic synapses and subject to biologically-inspired forms of learning can now perform feats of prediction and control previously thought to be the sole purview of the human brain, and increasingly permeate all aspects of digital life (Hassabis et al., 2017; Krizhevsky et al., 2012; Levine et al., 2018). Neurorobotics - the study of robots controlled by artificial nervous systems - leverages much of this synergy and is proving a powerful method for developing and validating computational models of brain function (Krichmar et al., 2018; Falotico et al., 2017).

Neurorobotics also has enormous and largely untapped potential as a neuroscience education technology because it combines multiple activities with clear educational benefits. (1) Robotics is a highly motivating and effective framework for teaching STEM in schools (Barker, 2012; Benitti, 2012; Karim et al., 2015), including to underrepresented students (Ludi, 2012; Rosen et al., 2012; Yuen et al., 2013; Weinberg et al. 2007). (2) The process of designing, testing and modifying neurorobot brains with interesting behavioral and psychological capacities engages students in active learning, which has been shown to improve STEM outcomes (Freeman et al., 2014), especially among disadvantaged students (Cervantes et al., 2015; Haak et al., 2011; Kanter & Konstantopoulos, 2010). (3) Finally, neurorobotics combines robotics and active learning with neuroscience, a highly multidisciplinary subject that presents itself in a wide array of real-life situations and readily appeals to the public (Sperduti et al., 2012; Frazzetto & Anker, 2009). The aim of neurorobotics is convincing robotic embodiment of attention, emotion, decision-making and many other mental capacities that are inherently interesting to students. Given user-friendly and affordable robot hardware, intuitive brain design and visualization software, and well-researched curriculum, educational neurorobotics has the potential to revolutionize neuroscience tuition, STEM education and the understanding of the brain.

Educational neurorobotics is a small but growing area of research and development. Iguana Robotics developed perhaps the first neurorobot for education - an inexpensive four-legged robot that uses capacitors and resistors to emulate neural networks and move (Lewis & Rogers, 2005). Middle and high-school students were readily motivated to modify these neural networks in order to change the robot’s gait, and demonstrated improved neuroscience attitudes as a result. NeuroTinker has more recently developed LED-equipped hardware modules that emulate individual neurons and can be connected into small neural circuits and attached to sensors and motors. Undergraduate students demonstrated improved understanding of neuroscience concepts after using the modules (Petto et al., 2017). Robert Calin-Jageman’s Cartoon Network is an educational neural network simulator that can connect via USB to the Finch Robot (BirdBrain Technologies LLC), a mobile robot with temperature-, light- and touch sensors, motorized wheels, lights and buzzers. Cartoon Network allows students to use different types of neurons and synapses to build neural circuits and control the Finch Robot, and generated promising results in workshops with undergraduates and teachers (Calin-Jageman, 2017; Calin-Jageman, 2018). Asaph Zylbertal’s NeuronCAD is a Raspberry Pi-based neurorobot that uses simulated neurons to control motors and process input from a camera (Zylbertal, 2016) but the project’s educational aims have not yet been implemented. Finally, Martin Sanchez at University Pompeu Fabra organizes an annual educational neurorobotics project as part of the Barcelona International Youth Science Challenge (Sanchez, n.d.). We set out to expand on these promising developments in educational neurorobotics and extend the range of brain functions and biologically-inspired neural networks students are able to create and the ways in which these networks can be visualized, analyzed and modified.

We have developed a neurorobot for high school neuroscience education that combines easy-to-use brain simulation and brain design software with affordable, wireless, camera-equipped do-it-yourself (DIY) hardware. Our DIY neurorobot is a mobile robot that uses wheels, a camera, a speaker and a distance sensor to navigate and interact with its environment. To keep hardware cost low while allowing students to leverage compute-intensive graphical user interface, brain simulation and machine learning functionality we chose to perform most computations on a wirelessly connected laptop that receives sensory input from the robot, extracts sensory features, simulates user-defined Izhikevich-type neural networks (Izhikevich, 2003) and sends commands back to the robot’s motors and speaker in real-time (Figure 1). The neurorobot consists of generic hardware components that can be purchased online at a total cost of about $150 and assembled in under 4 hrs with a soldering iron and a glue gun. A parts list and assembly instructions are provided below.

**Figure 1.**
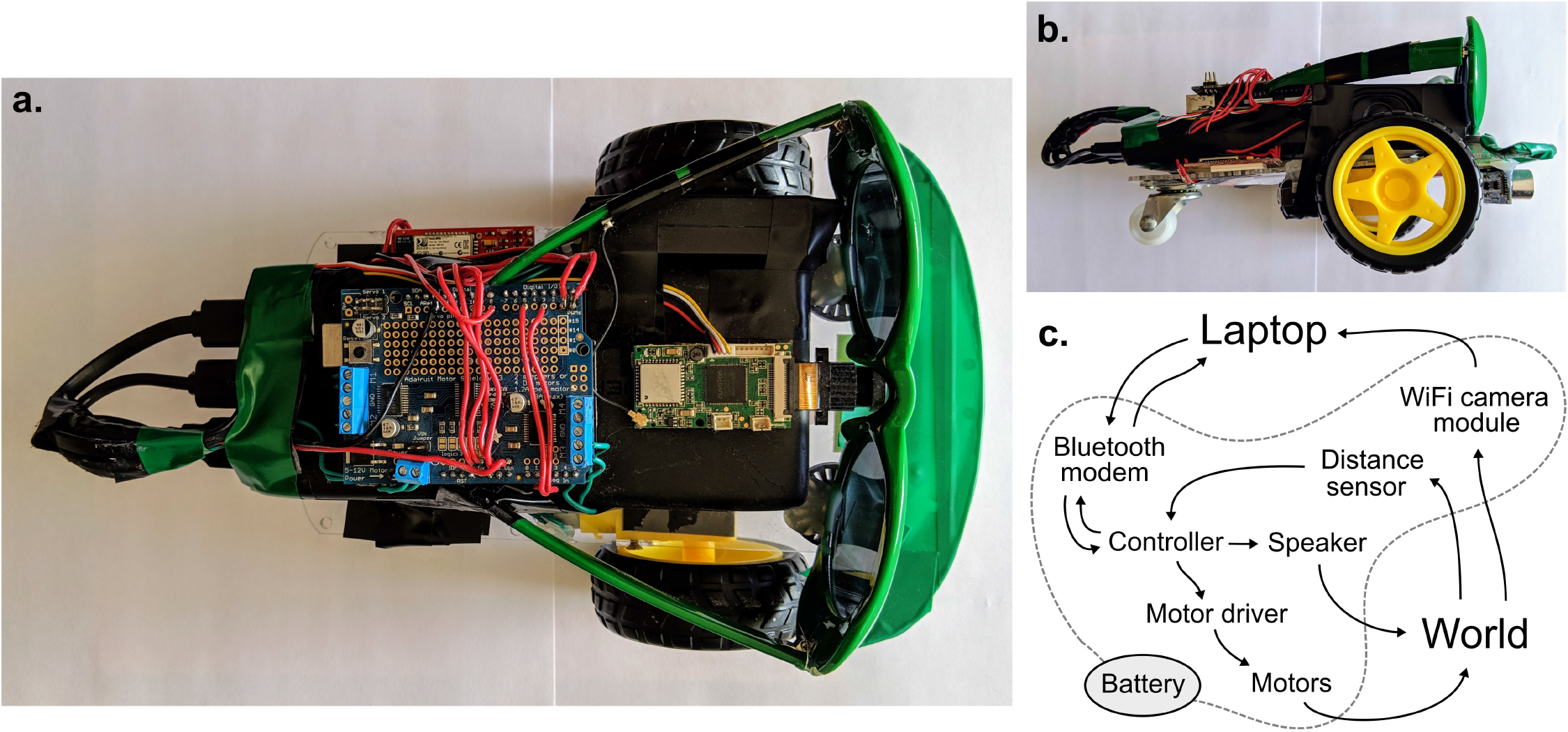
A DIY neurorobot for education. **a-b**. Top and sideways views showing the neurorobot’s chassis, battery, controller(with motor shield on top), WiFi camera module, speaker, distance sensor, bluetooth modem and sunglasses. **c.** Schematic showing the flow of signals in the system.

We have also developed an application that performs real-time visualization of the neurorobot’s brain and visual input, and allows user-controlled delivery of dopamine-like rewards and other commands (Figure 2). The app includes a brain design environment for building neural networks, either neuron-by-neuron and synapse-by-synapse or by algorithmic definition of larger networks. The Neurorobot App is written in Matlab and is available to download at github.com/backyardbrains/neurorobot. Although we recommend using neurorobot hardware, the app can run without it, and can use a normal webcamera to acquire visual input.

**Figure 2.**
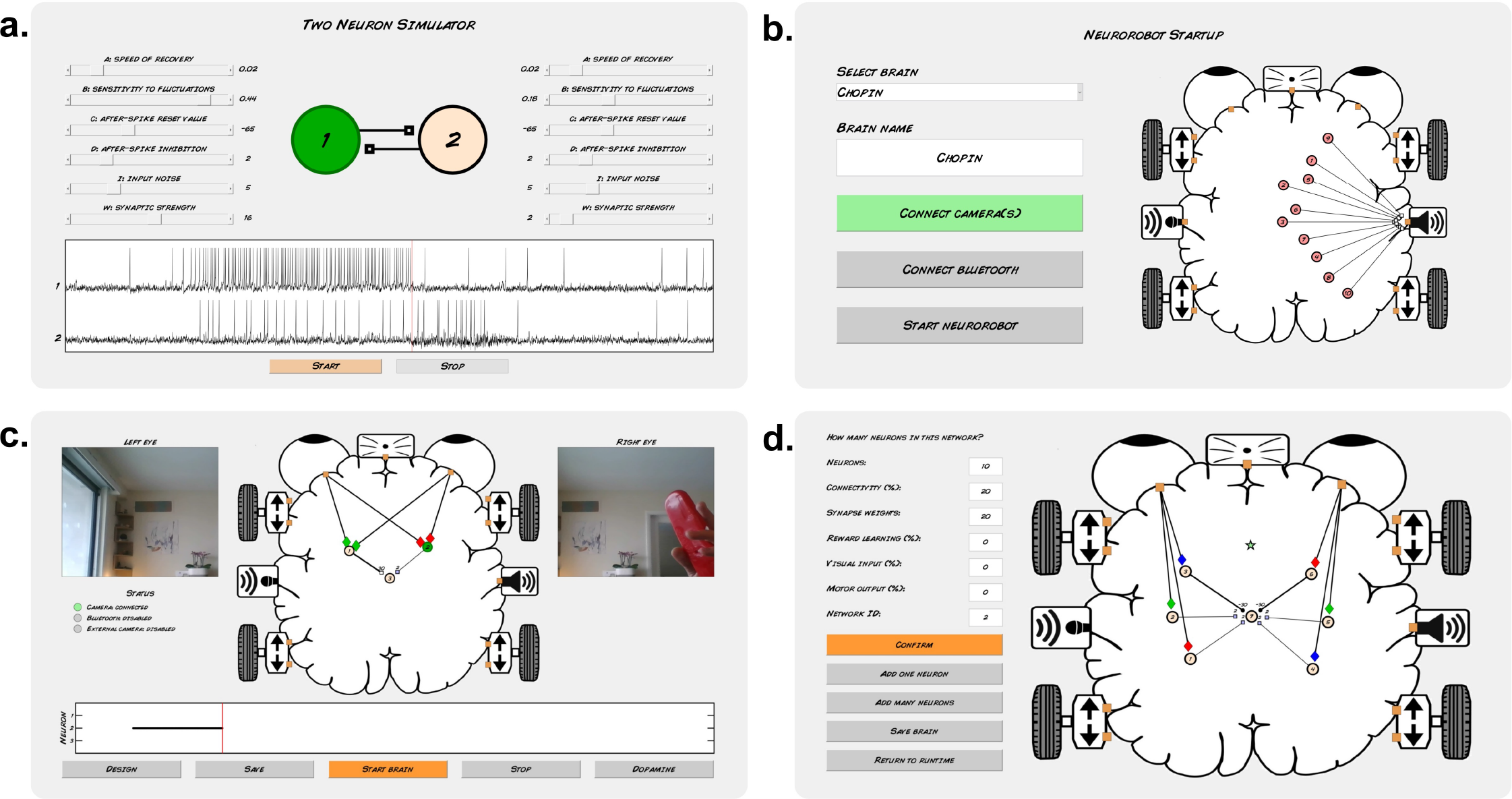
Neurorobot App. **a.** Two Neuron Simulator. **b.** Startup mode. **c.** Runtime mode. **d.** Brain design mode

To provide an initial assessment of the educational value of our neurorobot, we developed an introductory Neurorobot Workshop for high school students (Figure 3). In this workshop, students are first familiarized with the behavior of the Izhikevich neuron model and shown how to connect such neurons with synapses to produce goal-directed behavior (Braitenberg, 1986). Students then investigate Hebbian learning as they train their neurorobot to remember new visual stimuli. Finally, students explore action selection and reinforcement learning by using a “dopamine button” to teach their robot how to behave in different sensory contexts. We tested the Neurorobot Workshop with high school students (n = 9) and found that students were able to complete all exercises in under 3 hrs. In a post-workshop survey, students reported having gained the ability to work with computational models of neurons and synapses, and to develop neural networks that perform specific functions, including goal-directed behavior and memory (Figure 4).

**Figure 3.**
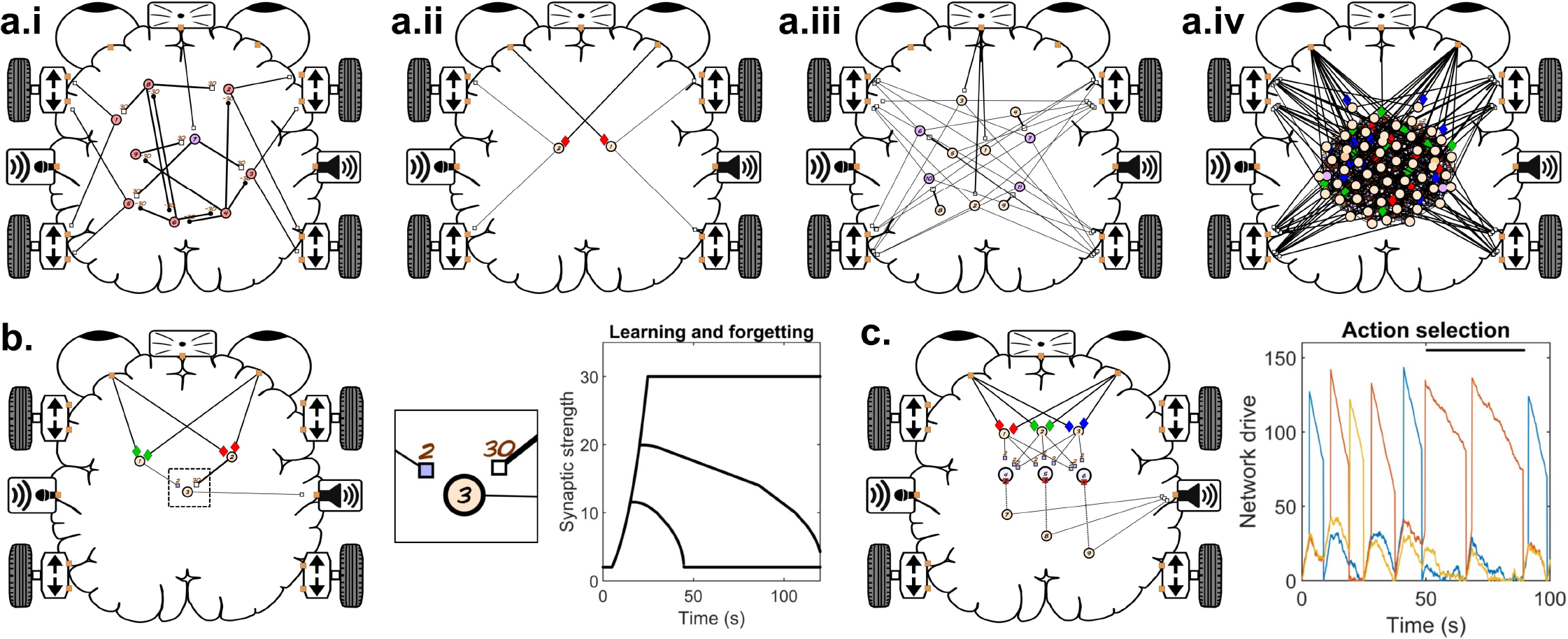
Neurorobot Workshop. **a.** Brains used to investigate spontaneous behavior (**a.i**), directed navigation (Braitenberg vehicle, **a.ii**), slow exploration (**a.iii**) and random networks (**a.iv**). In a Braitenberg vehicle, detection of a visual feature in either half of the visual field activates motors on the opposite side of the robot, allowing it to approach a target even if the target moves around. **b.** Brain used to investigate Hebbian learning. Plot shows synaptic decay rates for 3 durations of training. **c.** Brain used to investigate adaptive action selection. Plot shows network drive for 3 networks during normal and rewarded (black bar) behavior.

**Figure 4.**
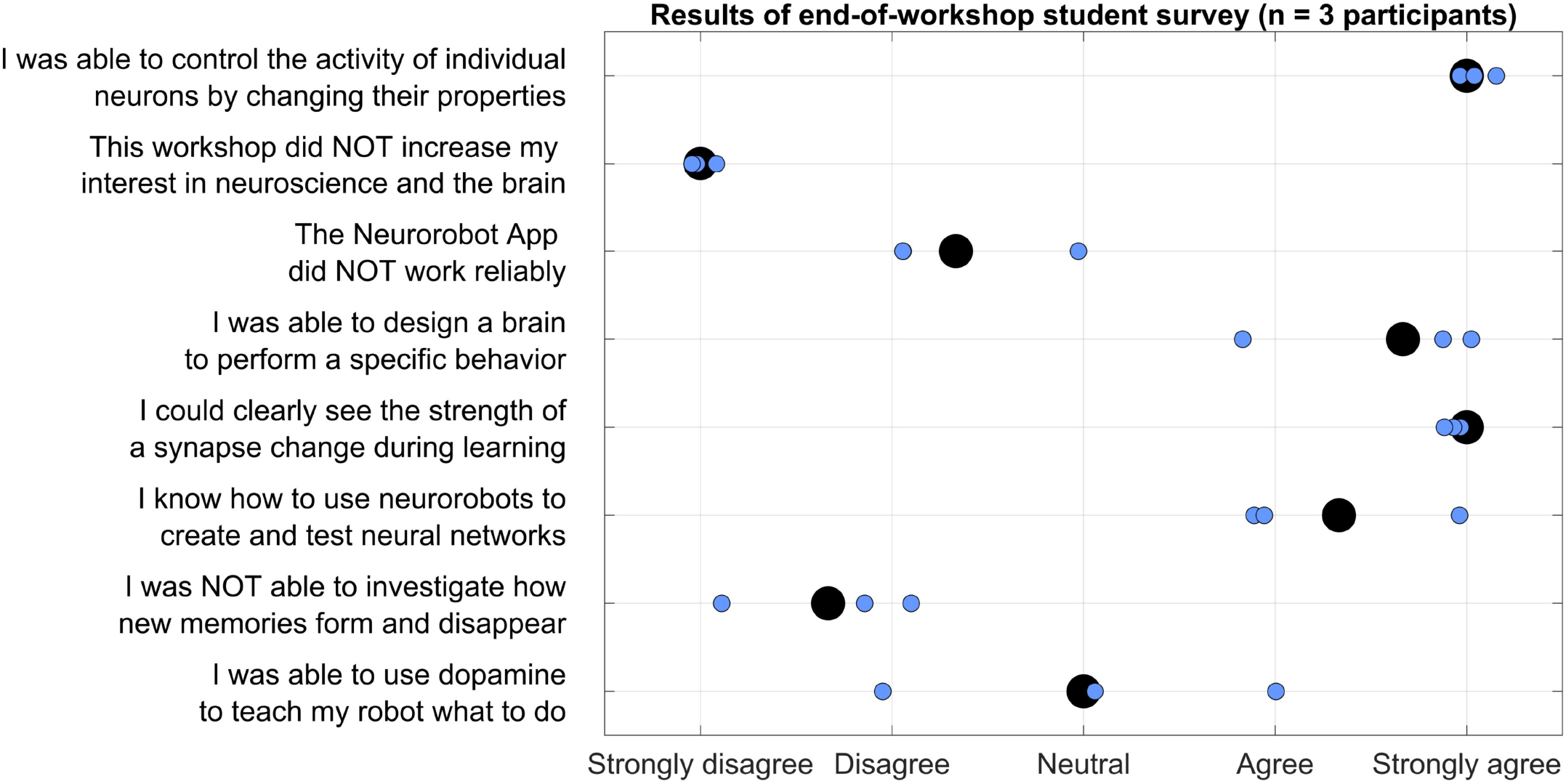
Survey results. Results of an end-of-workshop survey completed by 3 students. Black dots indicate average responses.

## 2. METHODS

### 2a. Neurorobot hardware

We developed our DIY neurorobot hardware design (Figure 1) with the aim of making neurorobotics accessible and appealing to a wide range of learners and learning communities. To accomplish this, we sought to use affordable components that can be purchased online and easily assembled, while at the same time allowing fast, wireless robot mobility, real-time processing of 720p video, audio communication, and simulation of relatively large spiking neural networks. The neurorobot uses a plastic chassis with two motorized wheels and a single swivel wheel. Double-sided tape was used to attach a 22000 mAh battery with 3 USB ports to the chassis. An UNO R3 controller with an Adafruit motor shield was affixed on top of the battery. A RAK5206 WiFi camera module was also attached on top of the battery with the camera attached to the forward-facing side of the battery. An 8 ohm speaker was taped to the side of the battery. A HC-SR04 ultrasonic distance sensor was attached to the forward-facing underside of the chassis with a glue gun. A SparkFun BlueSMiRF bluetooth modem was attached to the chassis, next to the battery. Finally, to increase the robot’s appeal to high school students, a pair of sunglasses was attached and color-matched tape was used to decorate the front of the chassis. A wiring diagram and step-by-step assembly instructions are provided in the Supplemental Information.

The neurorobot’s UNO R3 controller firmware is written in C/C++ and is available to download at github.com/backyardbrains/neurorobot. The controller communicates via the bluetooth modem with the Neurorobot App, which runs on a dedicated laptop (Figure 1c). Every ~100 ms the controller looks for a 5 byte package from the laptop representing a speed (0-250) and direction (1 = forward, 2 = backward) for the left motor (bytes 1-2), a speed and direction for the right motor (bytes 3-4) and an output frequency for the speaker (31-4978 Hz, 8-bit resolution). If the package is available, motor and speaker states are updated accordingly. In the same 10 Hz cycle, the controller also uses the ultrasonic sensor to estimate the distance to the nearest object in front of the neurorobot and sends this distance to the laptop (range: 4-300 cm, 32-bit resolution). In parallel, the neurorobot’s RAK5206 WiFi camera module collects 720p color images at 10 frames per second and sends them via WiFi to the dedicated laptop. We are currently developing a Matlab/C++ library that will allow the WiFi camera module to perform all wireless communication, obviating the need for the bluetooth modem.

### 2b. Neurorobot App

We developed a Matlab-based app to enable students with no background in neuroscience or programming to design spiking neural networks for the DIY neurorobot, and to visualize and interact with those neural networks in real-time. We chose to implement the Izhikevich neuron (Izhikevich, 2003), a spiking neuron model that balances realistic-looking membrane potentials and spike patterns with relatively light compute load. The Neurorobot App (Figure 2) simulates neurons at a speed of 1000 Hz as required by the Izhikevich formalism, while performing all other processes including audiovisual processing, plasticity and brain visualization at a speed of 10 Hz. Synaptic connections between neurons can be either excitatory or inhibitory (measured in mV, range: −30 to 30 mV). Individual excitatory synapses can be made subject to Hebbian learning and will then be strengthened if their presynaptic neuron fires simultaneously with or just before their postsynaptic neuron. This synaptic reinforcement is subject to inverse exponential decay and disappears completely after a few minutes unless the synaptic reinforcement is sufficiently strong (>24 mV) to form long-term memory. Hebbian learning can furthermore be made conditional on simultaneous delivery of dopamine-like reward signal.

To help students familiarize themselves with Izhikevich neurons the Neurorobot App features the Two Neuron Simulator (Figure 2a), which simulates a pair of neurons connected by reciprocal excitatory synapses. Sliders update the neuronal activity parameters, input noise and synaptic strengths of both neurons in real-time. The Two Neuron Simulator can be launched from the Neurorobot App startup menu.

In addition to simulating neurons and their synaptic connections, the Neurorobot App optionally implements action selection functionality modeled on the dynamics of action selection in the basal ganglia (Grillner, 2006; Seth et al., 2012; Prescott et al., 2006; Bolado-Gomez & Gurney, 2013). Each neuron is assigned a network ID during the brain design process and each such network is associated with a stochastically increasing level of ‘drive’ which reflects the network’s likelihood of being selected by the basal ganglia. Only neurons with a selected network ID can fire spikes (i.e. control behavior). If the drive of a network crosses a threshold the network may be selected (the selection process is currently implemented algorithmically, not neuronally). If selected, a network’s drive is significantly increased. While selected, the network will inhibit the drive of all other networks but will also slowly lose its own drive unless excitatory synaptic inputs or dopamine-like reward signals are provided. A special ‘medium spiny’ neuron type can be used to receive sensory and other neuronal inputs intended to modulate the drive of a specific network in different contexts, mirroring corticostriatal input to the basal ganglia. Synaptic inputs onto a medium spiny neuron increase the drive of that neuron’s network proportionally. Network ID 1 is exempt from selection to allow for uninterrupted sensory and other neuronal activity.

In the Neurorobot App, neurons and their connections are represented within a brain-shaped workspace (Figure 2b-d). The workspace is lined with icons representing camera input, distance input, audio input (not yet in use), motor output and sound output. Neurons are represented within this workspace as colored circles connected by axons (black lines) that end in synapses (small rectangles or circles representing excitatory and inhibitory synapses, respectively).

The Neurorobot App has three modes of operation: startup, runtime and design. In startup mode (Figure 2b), students are able to load and preview different brains, and establish their robot’s WiFi and bluetooth connections. In runtime mode (Figure 2c), students can visualize their robot’s visual input, brain activity and brain structure in real-time, and send dopamine-like reward signals to the brain. Neuronal spikes are indicated by black markers in a continually updated raster plot, and by the spiking cell body briefly turning green. The width of each line representing an axon indicates the absolute strength of that synaptic connection and is also updated in real-time. In design mode (Figure 2d), students are able to add individual neurons to the brain by selecting from a range of pre-specified neuron types with different firing properties, and connect them to other neurons by means of excitatory or inhibitory synapses with different strengths and plasticity rules. Students can also assign neurons a range of sensory preferences (colors, objects, distances) and motor outputs (speed and direction of movement, sound frequencies) by drawing axonal connections between neurons and the various icons lining the brain-shaped workspace. Object recognition is accomplished using Matlab’s Deep Learning Toolbox and requires a relatively high-performance graphics card. However, colors provide sufficient sensory diversity for all exercises described in the Neurorobot Workshop below. Design mode also allows students to add groups of neurons to the brain by algorithmically defining their properties (Figure 2d). The code needed to run the Neurorobot App is available at github.com/backyardbrains/neurorobot and does not require neurorobot hardware to run (see Supplemental Information).

### 2c. Neurorobot Workshop

We designed and performed a short Neurorobot Workshop with the aim of introducing students to our neurorobot and providing an initial evaluation of its potential as a tool to teach computational neuroscience in high schools (Figure 3). Our protocol was approved by IntegReview IRB (March 21, 2018, protocol number 5552). The workshop was conducted on 3 occasions and involved a total of 9 high school students aged 13-17. Each workshop involved 2-4 students, many of which were current or former members of the highly qualified FTC Team 9794 Wizards.exe. Each student also had their own laptop and robot to work with. These ideal conditions allowed us to focus on usability and less on issues of comprehension and miscommunication that can arise in larger groups of less experienced students. None of the students were attending high schools with a dedicated neuroscience program or had other training in neuroscience. The workshop instructor (Dr. Harris) used a separate laptop and robot to perform demonstrations. 1-2 parents or teachers attended each workshop. Instructions for starting and using the neurorobot are provided in the Supplemental Information.

The workshop began with the instructor providing a short introduction to the concept of brain-based robots. Students then started the Neurorobot App by navigating to the app folder in Matlab and running the script *neurorobot.m*. Students began by accessing the Two Neuron Simulator (Figure 2a) and were taught to identify spikes, reduce input noise, increase or decrease spike rate, and trigger postsynaptic spikes by varying synaptic strength. The aim of this exercise was to familiarize students with spiking neurons, with the fact that neurons can be quiet or spontaneously active, and with the synaptic strength needed to reliably trigger spikes in a postsynaptic neuron.

Students were then instructed to use the Neurorobot App to design a brain that would move forward in response to seeing a specific color. (This can be accomplished by adding a single neuron to the brain, assigning it a color preference, and extending axons to the forward-going motors on both sides of the robot.) This required students to learn how to transition between the startup, runtime and design modes of the app, to add a neuron of the correct (i.e. quiet) type to the workspace, and to assign the neuron specific sensory inputs and motor outputs. Students were then encouraged to modify the brain so that the robot would spin around in response to seeing a different color, move backward in response to the distance sensor registering a nearby object, and to produce distinct tones during each behavior. The aim of this exercise was to familiarize students with the Neurorobot App and help them develop an understanding of how functionally distinct neural circuits can co-exist in a single brain.

Students were then encouraged to explore the various pre-configured brains available from the startup menu (e.g. Figure 3a.i-iv), to try to understand how it is possible for many of them to generate distinct spontaneous behaviors in the absence of specific sensory inputs, and to incorporate some of the operative neural mechanisms in their own brain designs. This required students to learn how to load different brains and analyse them in order to identify neuronal properties that contribute to spontaneous behavior (e.g. spontaneously active or bursting neurons). Students were also introduced to the concept of a goal-directed Braitenberg vehicle (Braitenberg, 1986; Figure 3a.ii) and to random neural networks generated by defining neuronal and network properties probabilistically (Figure 3a.iv).

In the second (and shorter) part of the workshop students were first introduced to Hebbian learning and synaptic plasticity. Student were instructed to load a pre-configured brain consisting of three neurons (‘Betsy’, Figure 3b). Neuron 1 was responsive to green and projected a weak but plastic synapse to neuron 3. Neuron 2 was responsive to red and projected a strong but non-plastic synapse to neuron 3. Neuron 3 produced a sound output. Thus, showing the robot a red object led to activation of neurons 2 and 3 and the sound output, whereas showing the robot a green object only activated neuron 1. The challenge was to train the neurorobot to produce a sound in response to seeing the green color alone. The instructor explained the concept of Hebbian learning and showed students how to reinforce the synapse connecting neurons 1 and 3 by presenting the robot with both red and green colors simultaneously. Students were then asked to plot the strength of the plastic synapse in order to quantify how the duration of training (simultaneous stimulus presentation) affected the rate of learning and subsequent forgetting. The aim of the exercise was familiarize students with Hebbian synaptic reinforcement, which depends on simultaneous activation of a pre- and a postsynaptic neuron, to teach them how to read and plot synaptic strengths, and to introduce the idea that sufficient synaptic reinforcement can trigger long-term memory.

Students were then introduced to the concepts of action selection and reinforcement learning (Figure 3c). The instructor explained how the basal ganglia enables action selection in the vertebrate brain by selective disinhibition of specific behavior-generating neural networks (Grillner, 2006; Seth et al., 2012; Prescott et al., 2006; Bolado-Gomez & Gurney, 2013), and how this process is modulated by dopamine to promote behaviors that lead to reward. Students were shown how this selection process is implemented in the Neurorobot App by means of neural network IDs, thresholded levels of network drive and the medium spiny neuron type (see Section 2b). Students were then instructed to load a preconfigured brain that produces three different behaviors and can be conditioned using the dopamine button (‘Merlin’, Figure 3c). Students then trained the brain to perform one of its three behaviors in response to seeing a specific color by showing the color to the robot, waiting for it to perform the desired behavior, and then using the dopamine button to reinforce the color-behavior association.

At the end of the workshop, students completed an 8-question Likert-scale questionnaire designed to evaluate the educational benefits of the workshop (Figure 4). The second part of the workshop was only enacted at the last workshop event (n = 3 students), and only students who participated in both parts of the workshop were asked to complete the survey.

## 3. RESULTS

### 3a. App speed and performance

We perceived the stability and speed of the Neurorobot App to be critical to student engagement during the workshop. Early iterations of the Neurorobot App suffered from regular WiFi or bluetooth connection failures (“crashes”) that often required a system restart and irritated students. The current version of the app is more stable. When crashes do occur the app reconnects automatically in 6-7 seconds. Crashes are typically associated with trying to run 4 or more robots in close proximity, but the room and building in which the workshop takes place are also a factor. In the average workshop involving 5 simultaneously running neurorobots, 4 will perform well, with only one or two suffering occasional crashes (approx. 4-5 per hour).

The rendering speed of the Neurorobot App depends on the laptop used to run it. We found that laptops with 1.1 GHz dual-core CPUs, 8GB RAM and internal graphics (4165 MB total memory, 128 MB VRAM) were not able to render the app at an acceptable rate. The rendering of buttons and execution of their associated functions in brain design mode were particularly affected, resulting in Matlab errors and crashes as students attempted to start new processes before previously triggered ones had completed. We found that laptops with 2.80 GHz quad-core CPUs, 16GB RAM and NVIDIA GeForce GTX 1060 graphics (11156 MB total memory, 2987 MB VRAM) *were* able to render the app with only occasional delays. We used laptops with these specs, running Windows 10 and Matlab 2018a, in the workshops discussed here.

The neurorobot app is configured to run at 10 Hz, with its Izhikevich neurons running at 1000 Hz. With these settings we were able to simulate brains of up to 200 neurons, more than 10 times the number of neurons needed to run the workshop exercises. Nevertheless, improving our code to allow real-time simulation of much larger brains is a priority. We found that we had to use a slower rendering speed of 5 Hz when simulating larger brains or when using the Deep Learning Toolbox to perform object recognition.

The second persistent problem we encountered during workshops was color detection. In rooms with plenty of natural light our neurorobots were easily able to recognize and distinguish between red, green and blue objects. However, rooms with limited or amber ceiling lights were difficult to work in, with detection of green and blue being particularly affected. In one case this problem was so severe that the workshop had to be interrupted and the students relocated to a different room. We are currently working to improve the color detection functionality of the Neurorobot App, and recommend testing a range of colored objects in the same room and lighting conditions in which work with the app is to take place.

Two features of the brain design mode were not intuitive to students. First, the app assumed that the presynaptic origin of a synapse would be selected before its postsynaptic target. This was a natural way of establishing directed connections between pairs of neurons. However, it also meant that to make a neuron responsive to sensory input, the sensory input icon had to be selected before the target neuron. Similarly, to enable a neuron to produce motor or speaker output, the neuron had to be selected before the output icon. Although these requirements may reinforce the concept of directed signal flow in neural networks, students found the constraints frustrating. Students also found the process of modifying or deleting synapses confusing. To create a synapse, students had to extend an axon from the presynaptic origin (e.g. a neuron) to the postsynaptic target (e.g. another neuron). However, modifying or deleting an existing synapse also required students to extend an axon from the presynaptic origin to the postsynaptic target. Students repeatedly voiced the opinion that clicking directly on an existing axon or synapse to edit its properties would be more intuitive. We are currently working to solve both these user interface issues.

### 3b. Survey responses

Students’ responses to the statement “The Neurorobot App did NOT work reliably” ranged from disagree to neutral. This reflects the fact that the app is still at a prototype stage and subject to the “bugs” discussed above. We are exploring several approaches to improve app performance, including open-sourcing our code and implementing multi-core functionality.

Students strongly agreed with the statement “I was able to control the activity of individual neurons by changing their properties”. This indicates that the use of the Two Neuron Simulator at the start of the workshop and subsequent encouragement of students changing the properties of neurons from quiet to spontaneously active or vice versa during the workshop was successful. Students also agreed or strongly agreed with the statements “I was able to design a brain to perform a specific behavior” and “I know how to use neurorobots to create and test neural networks”. This reflects the fact that all students were able to complete the first part of the workshop in which they designed a brain to perform different sounds and movements in response to seeing different colors. This exercise typically elicited many positive exclamations as students saw their robots move for the first time.

Students strongly agreed with the statement “I could clearly see the strength of a synapse change during learning”, confirming that students learned how to observe this core feature of memory formation. Students also disagreed or strongly disagreed with the statement “I was NOT able to investigate how new memories form and disappear”. This reflects the fact that all students who participated in the second half of the workshop were able to train the brain Betsy to express a new color-behavior association.

Students’ responses to the statement “I was able to use dopamine to teach my robot what to do” ranged from disagree to agree. This final part of the workshop is clearly the most conceptually demanding, as it requires students to understand the idea of ongoing selection between competing neural networks and the modulation of this competition by sensory input and reward. To improve students’ understanding of this part of the workshop we have made three improvements: 1) neurons with the same network ID are now connected by dashed lines to indicate network membership; 2) medium spiny neurons are now larger than other neurons to indicate that they work differently; and 3) the color of medium spiny neurons now indicates their network’s level of drive (see Section 2b). We are also working to improve our presentation of this complex material.

Finally, all students strongly disagreed with the statement “This workshop did NOT increase my interest in neuroscience and the brain”, indicating that the workshop has a positive effect on students’ attitude towards neuroscience. Two students also indicated in the open “Ideas for improvement” section of the survey that more background information about neurons and synapses, as well as a detailed workshop program, would have been helpful.

## 4. DISCUSSION

Neurorobotics offer students a unique opportunity to learn neuroscience and computational methods by building and interacting with embodied models of neurons and brains. To promote neurorobot-based neuroscience tuition in schools and educational neurorobotics as an area of research, we have provided here the instructions to build our DIY neurorobot, the Matlab-code of the associated Neurorobot App, and the contents and results of a Neurorobot Workshop for high school students. We found that students enjoyed the Neurorobot Workshop, were generally able to understand and complete the exercises we presented, and expressed competence and confidence in computational neuroscience in a post-workshop survey.

Development of our educational neurorobotics platform is ongoing. In the near future we will remove the need for the bluetooth modem by conducting all serial communication via WiFi. This involves creating a new library for communication between Matlab and the RAK5206 WiFi module, as the HebiCam library we are currently using for this purpose only allows video transmission. We are also working on a lighter, fabricated hardware design with microphone, gyroscope and accelerometer input, motor encoders and multi-color LEDs.

On the software side there are numerous near-term improvements that would improve usability, including tools to edit neurons and synapses en masse, and methods for working with larger brains (e.g. zooming in and out, hiding different types of brain structure). An efficient search of the space of possible neural networks with the aim of discovering novel, engaging behavioral outputs would allow students to more quickly create and train interesting brains in the classroom. Faster and richer sensory feature extraction would allow students to design brains that can recognize and adapt to key features of their local environment. Perhaps the most important near-term goal is to expand the library of brains available to students, to include interesting, useful, well-understood features such as retinotopy and place cells, and to develop creative pedagogic exercises to introduce these brains to students. We are particularly interested in creating brains that make use of the dopaminergic neuron type, activation of which generates reward signal and enables reinforcement learning without the need for the dopamine button.

As researchers and educators exploring the still nascent field of educational neurorobotics we are faced with numerous interesting questions. What types of sensory features are most useful to students and how should they be made accessible in the Neurorobot App? What types of neurons and neural circuits do students need to be able to deploy with a single click? What types of exercises, brains and behaviors do students prefer to work with, and why? How should existing neurorobotics research and computational brain models be translated into forms suitable for the K-12 classroom? How should educational neurorobotics be combined with project-based learning? How can we make teachers confident about using neurorobots to teach neuroscience? And how should neuromorphic hardware be incorporated into educational neurorobots? It's an exciting time to bring neurorobots to the classroom!

## ACKNOWLEDGEMENTS

Research reported in this publication was supported by a Small Business Innovation Research grant #1R43NS108850 from the National Institute of Neurological Disorders and Stroke of the National Institutes of Health. CAH and GJG are Principal Investigators on this grant. GJG is a co-founder and co-owner of Backyard Brains, Inc., a company that sells the open-source hardware described in this paper.

## REFERENCES

Barker, B. S. (Ed.). (2012). Robots in K-12 Education: A New Technology for Learning: A New Technology for Learning. IGI Global.

Benitti, F. B. V. (2012). Exploring the educational potential of robotics in schools: A systematic review. Computers & Education, 58(3), 978–988.

Bolado-Gomez, R., & Gurney, K. (2013). A biologically plausible embodied model of action discovery. Frontiers in neurorobotics, 7, 4.

Braitenberg, V. (1986). Vehicles: Experiments in synthetic psychology. MIT press.

Calin-Jageman, R. J. (2017). Cartoon Network: A tool for open-ended exploration of neural circuits. Journal of Undergraduate Neuroscience Education, 16(1), A41.

Calin-Jageman, R. J. (2018). Cartoon Network Update: New Features for Exploring of Neural Circuits. Journal of Undergraduate Neuroscience Education, 16(3), A195.

Cervantes, B., Hemmer, L., & Kouzekanani, K. (2015). The Impact of Project-Based Learning on Minority Student Achievement: Implications for School Redesign. Education Leadership Review of Doctoral Research, 2(2), 50–66.

Dekker, S., & Jolles, J. (2015). Teaching about “brain and learning” in high school biology classes: Effects on teachers' knowledge and students' theory of intelligence. Frontiers in psychology, 6.

Falotico, E., Vannucci, L., Ambrosano, A., Albanese, U., Ulbrich, S., Vasquez Tieck, J. C., … & Cauli, N. (2017). Connecting artificial brains to robots in a comprehensive simulation framework: the neurorobotics platform. Frontiers in neurorobotics, 11, 2.

Frantz, K. J., McNerney, C. D., & Spitzer, N. C. (2009). We've got NERVE: a call to arms for neuroscience education. Journal of Neuroscience, 29(11), 3337–3339.

Frazzetto, G., & Anker, S. (2009). Neuroculture. Nature Reviews. Neuroscience, 10(11), 815.

Freeman, S., Eddy, S. L., McDonough, M., Smith, M. K., Okoroafor, N., Jordt, H., & Wenderoth, M. P. (2014). Active learning increases student performance in science, engineering, and mathematics. Proceedings of the National Academy of Sciences, 111(23), 8410–8415.

Fulop, R. M., & Tanner, K. D. (2012). Investigating high school students conceptualizations of the biological basis of learning. Advances in physiology education, 36(2), 131–142.

Gage, G. J. (2019) The Case for Neuroscience Research in the Classroom. Neuron, In press.

Grillner, S. (2006). Biological pattern generation: the cellular and computational logic of networks in motion. Neuron, 52(5), 751–766.

Haak, D. C., HilleRisLambers, J., Pitre, E., & Freeman, S. (2011). Increased structure and active learning reduce the achievement gap in introductory biology. Science, 332(6034), 1213–1216.

Hassabis, D., Kumaran, D., Summerfield, C., & Botvinick, M. (2017). Neuroscience-inspired artificial intelligence. Neuron, 95(2), 245–258.

Izhikevich, E. M. (2003). Simple model of spiking neurons. IEEE Transactions on neural networks, 14(6), 1569–1572.

Kanter, D. E., & Konstantopoulos, S. (2010). The impact of a project-based science curriculum on minority student achievement, attitudes, and careers: The effects of teacher content and pedagogical content knowledge and inquiry-based practices. Science Education, 94(5), 855–887.

Karim, M. E., Lemaignan, S., & Mondada, F. (2015, June). A review: Can robots reshape K-12 STEM education?. In Advanced Robotics and its Social Impacts (ARSO), 2015 IEEE International Workshop on (pp. 1–8). IEEE.

Krichmar, J. L. (2018). Neurorobotics—A Thriving Community and a Promising Pathway Toward Intelligent Cognitive Robots. Frontiers in Neurorobotics, 12.

Krizhevsky, A., Sutskever, I., & Hinton, G. E. (2012). Imagenet classification with deep convolutional neural networks. In Advances in neural information processing systems (pp. 1097–1105).

Labriole, M. (2010, July). Promoting brain-science literacy in the K-12 Classroom. In Cerebrum: the Dana forum on brain science (Vol. 2010). Dana Foundation.

Levine, S., Pastor, P., Krizhevsky, A., Ibarz, J., & Quillen, D. (2018). Learning hand-eye coordination for robotic grasping with deep learning and large-scale data collection. The International Journal of Robotics Research, 37(4-5), 421–436.

Lewis, A. & Rogers, J. J. (2005), SlugBug: A tool for neuroscience education developed at Iguana Robotics, Inc. http://www.iguana-robotics.com/publications/whitepaper.pdf. Accessed January 1, 2018.

Ludi, S. (2012). Educational robotics and broadening participation in STEM for underrepresented student groups. Robots in K-12 Education: A New Technology for Learning, 343–361.

Petto, A., Fredin, Z., & Burdo, J. (2017). The use of modular, electronic neuron simulators for neural circuit construction produces learning gains in an undergraduate anatomy and physiology course. Journal of Undergraduate Neuroscience Education, 15(2), A151.

Prescott, T. J., González, F. M. M., Gurney, K., Humphries, M. D., & Redgrave, P. (2006). A robot model of the basal ganglia: behavior and intrinsic processing. Neural Networks, 19(1), 31–61.

Rosen, J., Stillwell, F., & Usselman, M. (2012). Promoting diversity and public school success in robotics competitions. Robots in K-12 education: A new technology for learning, 326–342.

Sanchez, M., (n.d.), Neuro-robotics as a tool to understand the brain. https://biysc.org/programmes/research-projects/neuro-robotics-tool-understand-brain-0. Accessed March 29, 2019.

Seth, A. K., Prescott, T. J., & Bryson, J. J. (Eds.). (2011). Modelling natural action selection. Cambridge University Press.

Sperduti, A., Crivellaro, F., Rossi, P. F., & Bondioli, L. (2012). “Do Octopuses Have a Brain?” Knowledge, Perceptions and Attitudes towards Neuroscience at School. PloS one, 7(10), e47943.

Weinberg, J. B., Pettibone, J. C., Thomas, S. L., Stephen, M. L., & Stein, C. (2007, June). The impact of robot projects on girls’ attitudes toward science and engineering. In Workshop on Research in Robots for Education (Vol. 3, pp. 1-5).

Yuen, T. T., Ek, L. D., & Scheutze, A. (2013). Increasing participation from underrepresented minorities in STEM through robotics clubs. In Teaching, Assessment and Learning for Engineering (TALE), 2013 IEEE International Conference on (pp. 24–28). IEEE.

Zador, A. (2019). A Critique of Pure Learning: What Artificial Neural Networks can Learn from Animal Brains. bioRxiv DOI: 10.1101/582643.

Zylbertal, A. (2016). Neuron cad [video demonstration]. Retrieved January 1, 2018, from https://www.youtube.com/watch?v=vx5QH23PIk8.

