## Supplemental Information for "Short Neurorobotics Workshop for High School Students Promotes Competence and Confidence in Computational Neuroscience"

### Running the Neurorobot App without neurorobot hardware

To run the Neurorobot App without neurorobot hardware you need to change the variables *use_webcam* from 0 to 1, and *bluetooth_present* from 1 to 0. Both variables can be found at the beginning of *neurorobot.m*. If you haven't already, you will also need to install the Image Acquisition Toolbox Support Package for OS Generic Video Interface (available on the MathWorks website, link). Finally, you may need to change the settings of the *videoinput* function in *connect_rak.m* to recognize your webcamera (see *videoinput* documentation, link)

### Detailed assembly instructions

This section describes how to build one of our DIY neurorobots. In addition to the parts in the parts list below, the build requires a soldering iron, soldering wire, a glue gun, glue gun sticks, electrical wire, F/M jumper wire, insulating tape (black and one other color), double-sided mounting tape, 3 USB cables, small flat-head and Phillips-head screwdrivers and a pair of sunglasses. Those less experienced with soldering are advised to watch one of the many soldering tutorials available on YouTube before beginning.

**Parts list**

| Part | Approximate Cost | URL |
|---|---|---|
| Plastic chassis for mobile robots | $12 | link |
| 22000 mAh battery with 3 USB ports* | $40 | link |
| UNO R3 controller | $11 | link |
| Adafruit Motor Shield for Arduino V2 | $19 | link, link |
| RAK5206 WiFi camera module | $29 | link, link |
| 8 Ohm speaker | $8 | link |
| HC-SR04 distance sensor | $5 | link |
| SparkFun Bluetooth modem BlueSMiRF Silver | $27 | link, link |
| **Total** | **$152** | |

**This battery allows for many hours of continuous use and has been thoroughly tested by us. However, a smaller, lighter battery may be sufficient for most purposes, may cost less and may make the robot more agile.*

**Step-by-step instructions** (see also the wiring diagram below and Figure 1a-b)

1. **Chassis.** Strip the tips of four 8 cm lengths of wire. Solder the wires onto the contact pads of the two motors. Assemble the robot chassis (instructions are provided).
2. **Battery.** Use double-sided tape to attach the battery on top of the chassis. Place the battery so its ports are aligned with the back edge of the chassis. Cut the 3 USB cables so that each has 8 cm of cable attached. Strip the tip of each USB cable so that the internal voltage (red) and ground (black) wires are accessible. Strip the tip of six 8 cm lengths of wire and solder one onto each USB voltage or ground wire. Use black insulating tape to secure the cables. For additional instructions on how to modify USB cables, watch one of the many tutorials available on YouTube (e.g. link). Plug the USB cables into the

battery. Note: press and hold the battery power button to turn the battery on and off. Make sure the battery is turned off during assembly.

3. **Controller and motor shield.** Solder the headers and terminal blocks onto the motor shield (instructions provided on the Adafruit website, link). Attach the motor shield to the UNO R3 controller. Install the neurorobot firmware on your controller (available at github.com/backyardbrains/neurorobot). Also install the motor shield library (instructions provided on the Adafruit website, link). Use double-sided tape to attach the controller with motor shield on top of the battery so that its USB port is aligned with and doesn't cover the battery's four charge-indicating LEDs. The underside of the controller is uneven and may require 2-3 strips of tape to attach well to the battery. Solder the voltage and ground wires of one USB cable to the voltage input (Vin) and ground contacts of the controller. Attach the voltage and ground wires of another USB cable to the power terminal of the motor shield.
4. **Motor power.** Attach the wires from the left motor to the M1 terminal block and the wires from the right motor to the M3 terminal block (simply rearrange later if one or both motors are driving in the wrong direction).
5. **WiFi camera module.** Use double-sided tape to attach the RAK5206 module on top of the battery so that its camera can be attached with double-sided tape to the forward-facing side of the battery without stretching the camera wires. Attach the 4-contact cable (included with RAK5206) to pins 1-4 of the module. Strip the tips of the voltage (black) and ground (yellow) wires and attach them to the voltage and ground of the last USB cable.
6. **Bluetooth modem.** Strip the tips of four 8 cm lengths of wire. Solder the wires onto the TX-0, RX-1, VCC and GND contacts of the bluetooth modem. Use double-sided tape to attach the bluetooth modem on top of the chassis, to the left of the battery. Solder the TX-0 wire to pin 0 on the controller, the RX-1 wire to pin 1, the VCC wire to 3V and GND to ground. Be aware that the order of the pins of the bluetooth modem may differ from the wiring diagram below.
7. **Speaker.** Cut the speaker's wires at 8 cm and strip the tips. Use black insulating tape to attach the speaker to the left side of the battery so that the speaker's forward-facing side is aligned with the forward-facing side of the battery. Solder the speaker wires to (controller) pin 8 and ground. To use the speaker you must first create the pitches.h library on your controller (see the *tones* documentation on the Arduino website, link, link).
8. **Distance sensor.** Straighten the pins of the distance sensor if they are bent. Attach an F/M jumper wire to each pin. Cut and strip the tips of the jumper wires and solder on additional wire to extend the length of the jumper wires to 16 cm. Use double-sided tape to attach the distance sensor with wires to the underside of the chassis, facing forward. Use the glue gun to reinforce the attachment. Solder Vcc to 5V on the controller, Trig to pin 5, Echo to pin 3, and Gnd to ground. Use the glue gun to fasten wires to the underside of the chassis.
9. **Sunglasses.** Place the sunglasses so that the bridge straddles the camera without touching it. This may require first using the glue gun to attach two or more pieces of plastic (e.g. the currently unused black encoder wheels) to the top of the chassis, and using them as a platform to elevate the glasses. The temple tips of the glasses may also have to be cut to ensure a good fit. Use the glue gun to secure the underside of the glasses to the chassis/plastic platforms. Use tape to secure the temples of the glasses to the sides of the neurorobot.
10. **Antenna.** Firmly attach the WiFi antenna (included) to the RAK5206 module. Poor antenna attachment is a common cause of connection problems. Use black tape to secure the antenna to the left or right sunglass temple.
11. **Complete.** Use black tape to secure all wires. Use the glue gun to secure the wires of the motors, which otherwise tend to break, and any other wire that looks like it may come loose during repeated use. Use colored tape to decorate the front of the chassis, and to tie the USB cables together (see Figure 1a-b).

**Wiring diagram**

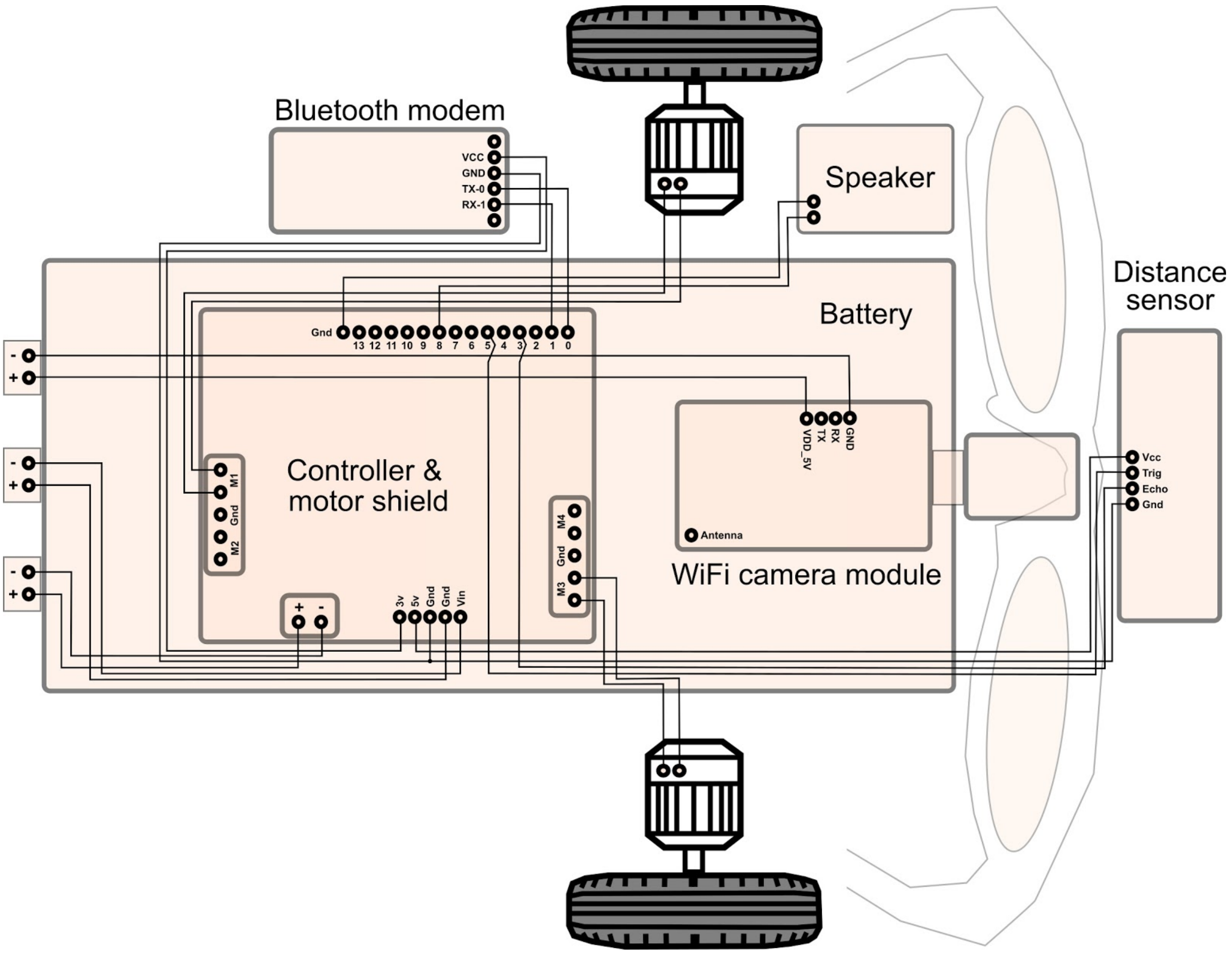


**Starting and using the neurorobot hardware**
Press and hold the battery power button to turn the robot on and off. After turning the robot on, wait a minute for the WiFi camera module's SSID to appear in your list of available WiFi networks - it will begin with the letters "LTH". Connect to it. The WiFi icon in the system tray will indicate "No network access" - this is expected and only indicates your lack of connection to the internet.

The first time you use the RAK5206 module you may need to change its settings to prevent connectivity problems in Matlab. To do this, make sure your computer is connected to the WiFi module's SSID, start Wisview for Windows (available at github.com/backyardbrains/neurorobot), press "Scan" to find the module and select the module's check box. Then, under "Set Video Param" change Video FPS to 10, Quality to 32, and Video GOP to 100. Click "Apply". A dialog box should appear confirming that the new settings have been applied to the module. Note: successfully connecting to RAK5206 by pressing "Connect camera" in startup mode will repeatedly generate the Matlab error message "Unexpected image dimensions. Skipping frame." for about 20 seconds - this can be ignored. We are currently working to remove this error message.

The first time you use the bluetooth modem you need to pair it with your computer. With the bluetooth modem turned on, open your computer's Bluetooth settings and find the bluetooth modem's ID - it will begin with the letters "RNBT". Pair with it. You may need to enter the password "1234". You also need to enter your modem's ID in the Neurorobot App. In *neurorobot.m*, find the variable *bluetooth_name* and make sure your bluetooth modem's ID is assigned to it. Connecting to the bluetooth modem by pressing "Connect bluetooth" in startup mode takes about 15 seconds.
